# Metabolic Reconstruction and Modeling Microbial Electrosynthesis

**DOI:** 10.1101/059410

**Authors:** Christopher W. Marshall, Daniel E. Ross, Kim M. Handley, Pamela B. Weisenhorn, Janaka N. Edirisinghe, Christopher S. Henry, Jack A. Gilbert, Harold D. May, R. Sean Norman

## Abstract

Microbial electrosynthesis is a renewable energy and chemical production platform that relies on microbial taxa to capture electrons from a cathode and fix carbon. Yet the metabolic capacity of multispecies microbial communities on electrosynthetic biocathodes remains unknown. We assembled 13 genomes from a high-performing electroacetogenic culture, and mapped their transcriptional activity from a range of conditions. This allowed us to create a metabolic model of the primary community members (*Acetobacterium, Sulfurospirillum*, and *Desulfovibrio*). *Acetobacterium* was the primary carbon fixer, and a keystone member of the community. Based on transcripts upregulated near the electrode surface, soluble hydrogenases and ferredoxins from *Acetobacterium* and hydrogenases, formate dehydrogenase, and cytochromes of *Desulfovibrio* were essential conduits for electron flow from the electrode into the electrosynthetic community. A nitrogenase gene cluster with an adjacent ferredoxin and one of two Rnf complexes within the genome of the *Acetobacterium* were also upregulated on the electrode. Nitrogenase is known to serve as a hydrogenase, thereby it would contribute to hydrogen production by the biocathode. Oxygenases of microaerobic members of the community throughout the cathode chamber, including *Sulfurospirillum* and *Rhodobacteraceae*, were expressed. While the reactors were maintained anaerobically, this gene expression would support anaerobic growth and thus electrosynthesis by scrubbing small amounts of O2 out of the reactor. These molecular discoveries and metabolic modeling now serve as a foundation for future examination and development of electrosynthetic microbial communities.

## Introduction

A microbial electrosynthesis system (MES) is a bioelectrochemical device that employs microbes to transport electrons from a cathode to protons and/or CO_2_. Thus, the microbes act as cathode catalysts to generate, or synthesize, a valuable product (Rabaey and Rozendal, 2010). This technology has immense potential, therefore understanding which microbes are capable of cathodic electron transfer and which metabolic pathways are involved in CO2 conversion to chemicals are critical to improving the performance of MESs. Furthermore, this work has broader environmental implications including understanding ecological aspects of one carbon metabolism and extracellular electron transfer relevant to global biogeochemical cycling. Technologically, these MESs could have a significant impact on geoengineering applications such as carbon capture and storage.

Advances in microbial electrosynthesis and related biocathode-driven processes primarily focus on the production of three compounds: hydrogen (electrohydrogenesis)(Rozendal *et al.*, 2008), methane (electromethanogenesis) (Cheng *et al.*, 2009), and acetate (electroacetogenesis) (Nevin *et al.*, 2010). However, microbial electrosynthesis can be used to generate alcohols and various short-chain fatty acids.\ (Steinbusch *et al.*, 2010). A thorough evaluation of a carbon dioxide fixing aerobic biocathode has also been reported (Wang *et al.*, 2015). To manipulate (and optimize) these systems we must understand the extracellular electron transfer (EET) enzymes or molecules involved in cathode oxidation, and we must determine how these cathodic electron transport components are coupled to energy conservation by the microbial cell. While an increasingly robust body of literature has been established relating to microbial communities metabolizing with an anodic electron acceptor (Ishii *et al.*, 2013, 2015; Kiely *et al.*, 2011), much is still unclear about the diverse metabolic capabilities of microorganisms and communities growing on a cathode (Tremblay and Zhang, 2015). We used metagenomics, metatranscriptomics, and metabolic flux modeling to elucidate carbon flux and EET mechanisms. We have deciphered the complex interactions between cooperating and competing members of the microbiome, producing opportunities to improve microbial electrosynthesis as well as understand multi-species biofilm communities.

## Materials and Methods

### Reactor design and operation

The reactors containing the microbial communities used for metagenomics and metatranscriptomics in this study were described in detail in two previous studies (Marshall *et al.*, 2012, 2013). Of the five MES reactors previously described, two reactors were selected for further analysis in this study. The first reactor, closed circuit (CC), was operated for 184 days before termination (MES-BW4 from previous manuscript is CC reactor here). The second reactor, open circuit (OC), was operated in closed circuit mode for 141 days before it was left at open circuit for three hours prior to termination, in order to control for biofilm effects of the closed circuit reactor (MES 1a from previous manuscript is OC reactor. The initial source of microorganisms was from a brewery wastewater retention basin that was then subjected to sequential transfers of the graphite and supernatant in bioelectrochemical systems before inoculation into the two MESs described in this study. The brewery wastewater inoculum was allowed to establish on graphite granule cathodes and then repeatedly washed with fresh defined medium containing per liter: 2.5 g NaHCO_3_, 0.6 g NaH_2_PO_4_ * H_2_O, 0.25 g NH_4_Cl, 0.212 g MgCl_2_, 0.1 g KCl, 0.03 g CaCl_2_, 20 mL vitamin solution, and 20 mL mineral solution. Subsequent, sludge-free transfers were made into 150 mL volume cathode chambers of MESs containing 30 g of graphite granules, 75 mL of medium, 80% N_2_ and 20% CO_2_ or 100% CO_2_ in the headspace, and poised at-590 mV vs. standard hydrogen electrode (SHE) for the duration of the experiments unless otherwise noted. The medium during the initial startup period did not contain sodium 2-bromoethanesulfonate (2-BES), but 10 mM 2-BES was periodically used to inhibit methanogenesis and maintain a predominantly acetogenic culture later in the study, including at the time of sampling for metagenome and metatranscriptome analyses (day 181). Upon termination of the experiment, supernatant and bulk granular cathodes were removed for DNA and RNA extraction. All samples were frozen in liquid nitrogen within 5 minutes of applied voltage interruption, except OC which was deliberately left at open circuit for 3 hours prior to freezing. A schematic of the experimental design can be found in Supplementary Figure 1.

**Figure 1.**
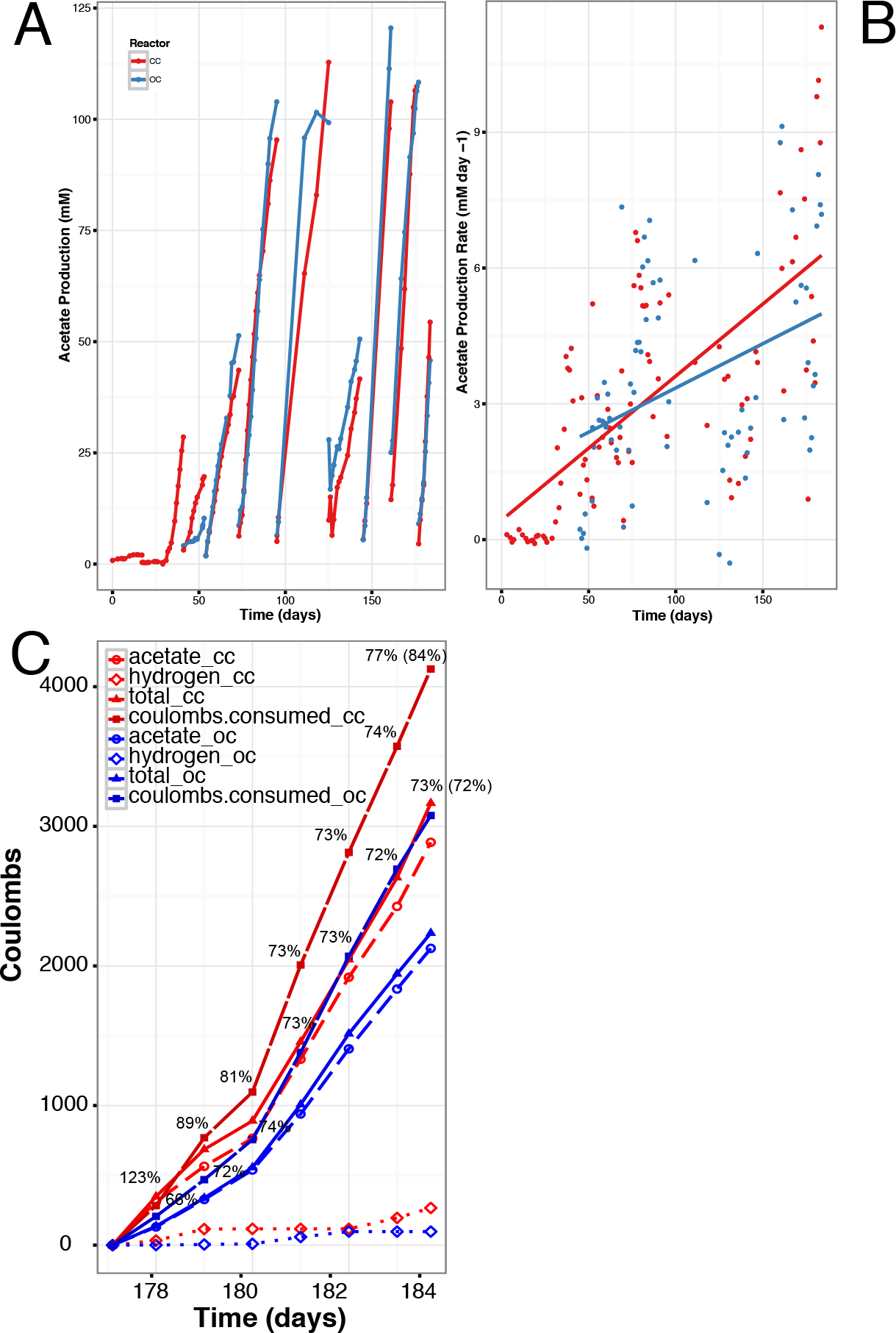
Reactor performance over time. (A) Acetate production in CC and OC-breaks in plotted lines indicate an exchange of the medium, (B) acetate production rate in CC and OC, and (C) accounting of the coulombs over a seven day period between medium exchanges in CC and OC, numbers are percent coulombic efficiency with total percent efficiency averaged over the 7 days in parentheses.

### DNA/RNA extraction and sequencing

DNA and RNA were processed as previously reported and is detailed in the Supplementary Online Methods. Whole genome shotgun sequencing of the cathode-associated microbial community from the closed circuit reactor (CCc) and the supernatant community from the closed circuit reactor (CCs) was accomplished with the Illumina MiSeq instrument yielding 250 bp paired-end reads totaling 9,793,154 reads with a total length of 2,325,061,111 bp for the attached microbial community, and 5,049,632 reads with a total length of 1,198,063,143 bp for the supernatant microbiome.

### Metagenome assembly and binning

Two metagenomes were assembled and annotated from the closed circuit reactor (CCc and CCs). Near full-length 16S rRNA sequences from the CCc and CCs metagenomes were assembled over 100 iterations of the Expectation-Maximization Iterative Reconstruction of Genes from the Environment (EMIRGE) method (Miller *et al.*, 2011), following initial read mapping to a modified version of the dereplicated version of the SILVA 108 small-subunit (SSU) rRNA database (Quast *et al.*, 2013). During iterations sequences 97% similar were clustered. Representative sequences were searched against the SSURef_111_NR_trunc database using BLASTN.

SPAdes (Bankevich *et al.*, 2012), Velvet+Metavelvet (Zerbino and Birney, 2008; Namiki *et al.*, 2012), Ray (Boisvert *et al.*, 2012), and IDBA-UD (Peng *et al.*, 2012) community genome assemblies were compared based on total length of the assembly, n50 score, and percent reads mapping back to the assemblies (Supplementary Table 1). Based on the assembly results, the SPAdes assembly was used to further analyze the metagenomes due to the highest percent of reads mapped to a threshold above 1kb (cathode: 95.95%, supernatant: 91.61%) and total assembly size (cathode: 67,511,580 bp above 1kb, supernatant: 42,119,276 bp). Following assembly, the SPAdes contigs were analyzed by Prodigal (Hyatt *et al.*, 2010) to predict protein-encoding genes. Prodigal-predicted amino acid sequences were then annotated using UBLAST searches (usearch64(Edgar, 2010)) against the uniref90 database (Suzek *et al.*, 2015) with an e-value cutoff of 100. These annotations, along with kmer coverage and GC content were used for preliminary genome binning. Emergent self-organizing maps (ESOM) were used to refine the metagenome bins based on tetranucleotide frequency and contig coverage on contigs greater than 4kb (Dick *et al.*, 2009). Smaller fragments (>1kb) were then projected onto the 4kb ESOM. To further refine the bins the multi-metagenome pipeline (Albertsen *et al.*, 2013) was used, leveraging differential genome coverages between CCc and CCs to aid contig identification. Results were compared to ESOM bins. Only contigs binned by both ESOM and multi-metagenome pipeline were used for reassemblies of individual genome bins. All reads associated with the contigs in each of the final bins were reassembled as single organism genomes by SPAdes using default options with the addition of the-careful flag and kmer sizes of 65, 77, 99, and 127. All contigs greater than 1 kb were uploaded to RAST (Aziz *et al.*, 2008) for final annotation using RASTtk (Brettin *et al.*, 2015). The re-assembled individual genome bins and one large “undetermined” bin were compiled into a combined metagenome for transcript mapping. See Supplementary Table 2 for bin summaries and completeness generated by CheckM (Parks *et al.*, 2015). The RASTtk annotations were compared to annotations (Wu *et al.*, 2011) from Pfam, COG, and a Blastx search of the non-redundant protein sequences database (Camacho *et al.*, 2009). The RAST-annotated protein encoding gene (peg) tags are used as identifiers throughout the manuscript with the bin abbreviation preceding the peg number. See Supplementary Table 3 for all annotations.

### Metatranscriptome

Four total metatranscriptomes were recovered from CCc, CCs, OCc, and OCs. The FASTX toolkit was used to filter the reads with a quality score lower than 30 and a length less than 50bp. Quality filtered reads from each sample were mapped to the CCc electrode metagenome contigs using BWA (Li and Durbin, 2009). Raw hit counts mapped to each RAST-annotated protein encoding gene in the combined metagenome were then used to calculate reads per kilobase gene length per millions of mapped reads (RPKM) values. RPKM values for each gene were also normalized to bin coverage based on single copy *recA* gene abundances (Supplementary Figure 6). This compensated for the abundance variations of the taxa between samples. Quantile normalization was used to evenly distribute the transcript values between sample conditions. Fold change for differential expression analysis between conditions was calculated by dividing RPKM/RecA-condition1 by RPKM/RecA-condition2. To eliminate 0 value errors and over interpretation of very low count genes, an empirically determined threshold value of 0.1 was added to all of the RPKM/RecA values.

### Metabolic model reconstruction, gapfilling, and curation

To construct models of our three highest relative abundance genomes *(Acetobacterium, Sulfurospirillum*, and *Desulfovibrio)*, we imported the RAST-annotated versions of these genomes into the DOE Systems Biology Knowledgebase (KBase). Once in KBase, we used the *Build Metabolic Model* app, which is based on the ModelSEED algorithm (Henry *et al.*, 2010), to construct one draft model for each genome. Once the draft models were complete, we curated the models based on literature data, KEGG (Kanehisa and Goto, 2000), and MetaCyc (Karp *et al.*, 2002)(Supplementary table 4c). This curation primarily involved the addition of electron utilization pathways *(Acetobacterium)*, addition of electron transport chain reactions *(Sulfurospirillum*, and *Desulfovibrio*), and adjustment of reaction directionality to prevent pathways from proceeding in unphysiological directions.

Next, we applied the *Gapfill metabolic model* KBase app to identify and fill all the gaps in the metabolic pathways of our models that might prevent them from producing biomass while operating in one of our hypothesized metabolic roles within the microbial community. In this gap-filling process, the model was augmented to include all the reactions contained in the ModelSEED database (available for download from GitHub (https://github.com/ModelSEED/ModelSEEDDatabase/tree/master/Biochemistry). Additionally, all reactions determined to be thermodynamically reversible (Henry *et al.*, 2006, 2009; Jankowski *et al.*, 2008) were adjusted to be reversible. Finally, flux balance analysis (FBA) (Orth *et al.*, 2010) was performed to generate a flux profile that produced biomass while minimizing the flux through all reactions and reaction directions that were not contained in the original model, consistent with previously published algorithms (Dreyfuss *et al.*, 2013; Latendresse, 2014). All reactions and reaction directions not included in the original model that had a nonzero flux in were then added to the model as the gap-filling solution. This subsequently permits the growth of the gap-filled model in the specified condition. *Acetobacterium* was gapfilled in a condition that reflected the only proposed metabolic role for this organism: direct consumption of electrons to fix CO2 coupled to the production of acetate. The *Sulfurospirillum* and *Desulfovibrio* models were gapfilled separately in two distinct conditions reflecting the potential roles proposed for these organisms: (i) autotrophic growth on CO_2_ and H_2_; or (ii) heterotrophic growth on acetate. This produced a total of four models: (i) autotrophic *Sulfurospirillum;* (ii) heterotrophic *Sulfurospirillum;* (iii) autotrophic *Desulfovibrio;* and (iv) heterotrophic *Desulfovibrio*. The detailed composition of the electrosynthetic, autotrophic, and heterotrophic media formulations used in the gapfilling analysis are included in the supplementary material (Supplementary Table 4).

All models were reconstructed, gapfilled, and curated in the KBase Narrative Interface, and the associated narratives include all parameters and media formulations used (https://narrative.kbase.us/narrative/ws.15248.obj.1). All model data is also available for download from the same narratives. The reactions for each model and condition can also be found in Supplementary Table 4a.

### Prediction of metabolic activity with comparison to expression data

Flux balance analysis (FBA) was applied with each of our gapfilled metabolic models to simulate the production of biomass on the same media formulations used in our gapfilling analysis (electrosynthetic, autotrophic, and heterotrophic media). Thus, we simulated the growth of our gapfilled *Acetobacterium* model in our electrosynthetic media; we simulated the growth of our autotrophic *Sulfurospirillum* and *Desulfovibrio* models in our autotrophic media; and we simulated the growth of our heterotrophic *Sulfurospirillum* and *Desulfovibrio* models in our heterotrophic media (all media formulations are listed in Supplementary table 4b). We then maximized the production of biomass in each model, subject to the constraints on nutrient uptake imposed by our media formulations, and once a solution was produced, we minimized the sum of the fluxes comprising the solution, in order to produce a single solution that is both simple and distinct.

We validated our flux solutions by comparing the active reactions in each flux profile against the actively expressed genes identified based on our transcriptomic data. In order to conduct this comparison, we first needed to classify the genes in our three species as either *active* or *inactive* based on the relative abundance of mapped RNA-seq reads. This process began by identifying an initial set of universal genes in each species that can be assumed to be *always active* (e.g. DNA polymerase subunits and tRNA synthetases). These genes are listed in Supplemental table 4d. We ranked these *always active* genes based on the abundance of reads mapped to them, identifying the number of reads mapped to the gene ranked at the 10^th^ percentile in this list as the threshold expression value. Then in the broader genome, we labeled any gene with mapped reads that exceeded the threshold as active, with all other genes being inactive. Finally, we calculated the binary distance between the transcriptome and flux profiles, excluding any reactions that had been gapfilled, to determine which predicted flux profiles yielded the greatest agreement (i.e. lower binary distance) with the transcriptomic data.

We carried out a comparative model analysis on thirty-three KEGG pathway categories involving gene presence and absence, transcriptomic data from CCc, flux analysis, and gap-filled reactions on each pathway category (Figure 3). Carbon fixation pathway categories (i.e. reductive TCA cycle and Wood-Ljungdahl pathways) were lumped into a single category called CO_2_ fixation. All of the metabolic models, their associated genomes, and the FBA analysis generated in this study are presented using the KBase Narrative Interface (NI) and are accessible at https://narrative.kbase.us/narrative/ws.15248.obj.1.

### Scanning electron microscopy-energy dispersive x-ray spectroscopy (SEM-EDX)

Cathodic graphite granules were fixed in 0.1 M sodium cacodylate buffer with 2% gluteraldehyde for three hours, washed in 2.5% osmium tetroxide, and then dehydrated with an ethanol dilution series using 0%, 25%, 50%, 75%, % and 100%. Samples were stored in a desiccator and then coated with Au and Pd using a Denton Vacuum sputter coater. Images were generated using a FEI Quanta 400 Scanning Electron Microscope.

### Data access

The metagenomic and metatranscriptome reads can be found in MG-RAST (Meyer *et al.*, 2008) under the ID numbers (and names) 4673464.3-4673467.3 (ARPA_metagenome) for the metagenome (http://metagenomics.anl.gov/linkin.cgi?project=15936) and 4536830.3-4536839.3 (ARPA_metatranscriptome) for the metatranscriptomes (http://metagenomics.anl.gov/linkin.cgi?project=6065). Assembled contigs for *Acetobacterium, Sulfurospirillum*, and *Desulfovibrio* can be accessed at https://narrative.kbase.us/narrative/ws.15248.obj.1. Raw Illumina sequencing reads have been deposited in the NCBI SRA database under Bioproject PRJNA245339. Additionally, sequencing and assembly files can be found online at https://github.com/sirmicrobe/electrosynthesis.

## Results

### Performance and compositions of microbial electrosynthesis systems

Two high performing MESs inoculated from an established electrosynthetic community were poised at-590 mV vs. a standard hydrogen electrode (SHE), and operated for over 100 days, generating an average percentage of total products formed of 65% acetate, 34% hydrogen, 0.4% formate, 0.3% propionate and 0.2% butyrate from CO_2_. At the conclusion of the experiment, one of the two reactors was left at open circuit for three hours to distinguish between transcripts influenced by the supply of electrons from the cathode compared to electrons supplied by hydrogen and/or other free metabolites in the biofilm. Similarly, metatranscriptome samples were taken from the electrode surface and compared to the supernatant. This led to four metatranscriptome samples from two reactors: closed circuit cathode (CCc), closed circuit supernatant (CCs), open circuit cathode (OCc), and open circuit supernatant (OCs). Coulombic efficiencies at time of sampling were 77% and 73% from CC and OC, respectively (Figure 1). Thirteen genome bins spanning 5 different phyla were recovered from the assembled metagenomes taken from the closed circuit MES (Figure 2). Near-complete genomes (89-100% complete) were produced for *Sulfurospirillum* str. MES7, *Acetobacterium* str. MES1, *Desulfovibrio* str. MES5, *Methanobacterium* str. MES13, *Bacteroides* str. MES9, *Geobacter* str. MES3, and *Sphaerochaeta* str. MES8 (Supplementary Table 2), and a second partially complete (~65%) *Desulfovibrio* str. MES6 genome was further obtained. *Acetobacterium, Sulfurospirillum*, and *Desulfovibrio* combined comprised 40-90% of the community in each condition, with a Rhodobacteraceae related organism also consistently represented (4-20% relative abundance) (Figure 2, Supplementary Figure 2). The 13 annotated genome bins recovered from the CC reactor were used as the basis for transcript mapping and analyses.

**Figure 2.**
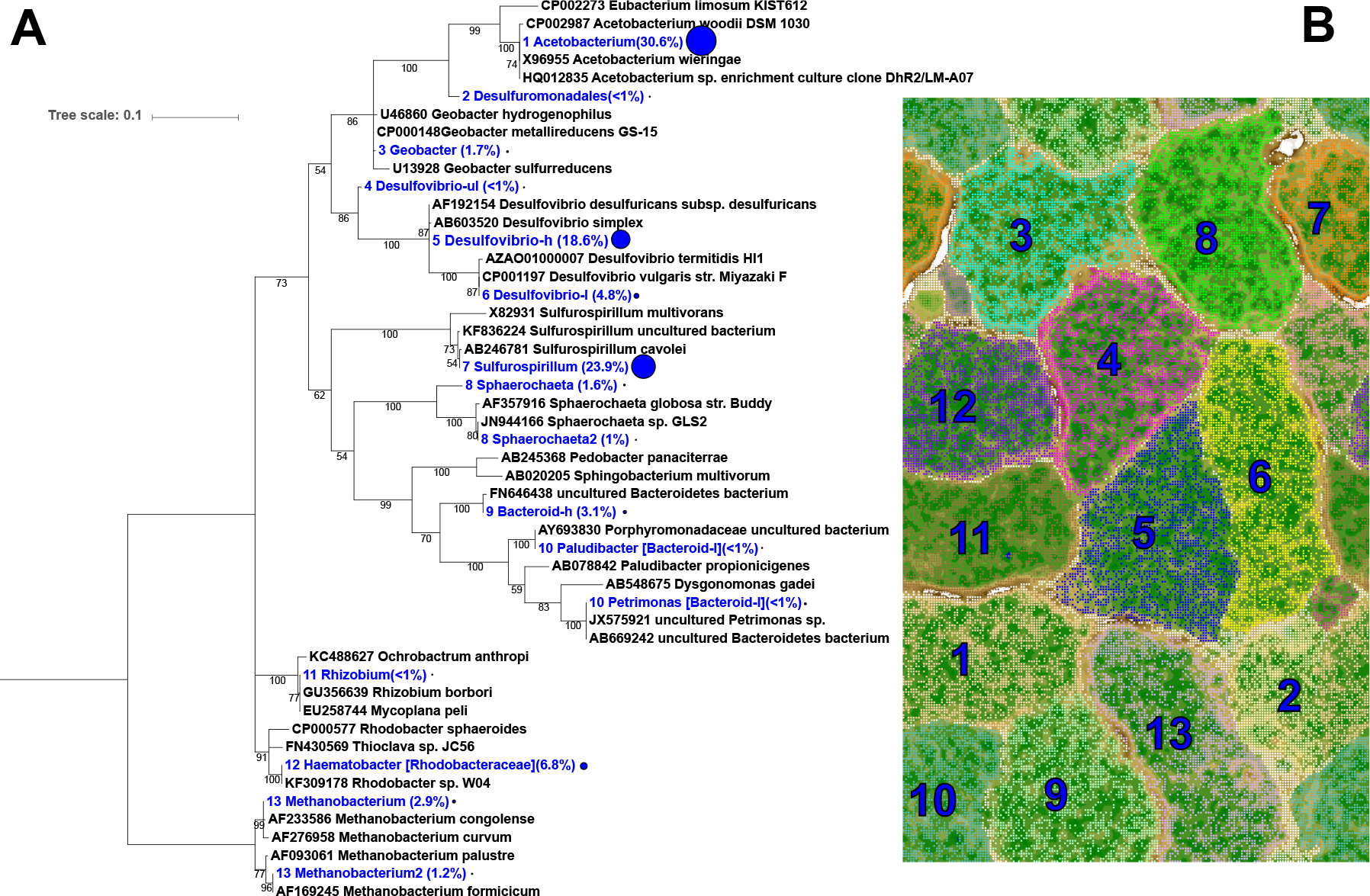
(A) Phylogenetic tree of the CCc microbial community using EMIRGE-based reconstructed 16S rRNA gene sequences (blue) and relative abundance values in parentheses and indicated by the relative size of the blue circle. Sequences were aligned using MUSCLE and the evolutionary history was inferred using the Maximum Likelihood method based on the Jukes-Cantor model. Bootstrapping support greater than 50% is indicated on the tree and is based on 1,000 iterations. (B) ESOM based on tetranucleotide frequency in the CC cathode metagenome.

### Description of the microbial catalysts

Three species, *Acetobacterium* str. MES1, *Sulfurospirillum* str. MES7, *Desulfovibrio* str. MES5, comprised 72% of the total abundance on the active cathode (CCc). The abundance, expression levels, and predominant activity in the reactor (acetogenesis) as well as the known physiology of similar organisms indicates that the *Acetobacterium* species is the main proponent of carbon fixation on the electrode and thus the keystone species in the microbial community. Genomic and transcriptomic evidence indicates the reconstructed *Acetobacterium* str. MES1 reduces CO_2_ to acetate through the Wood-Ljungdahl pathway (WLP). WLP-indicative acetyl-CoA synthase/carbon monoxide dehydrogenase (ACS/CODH, aceto.peg.1971-1981) and 5-methyltetrahydrofolate:corrinoid iron-sulfur protein methyltransferase (aceto.peg.1978) components were among the most highly expressed genes on the closed circuit electrode surface (Figure 3). Furthermore, the canonical enzyme ACS/CODH had on average >3-fold greater expression on the closed circuit electrode compared to the supernatant as well as higher expression on CCc compared to OCc, indicating that the electrode surface was the major source of acetate production. Interestingly, a *Clostridium-type* chain elongation pathway(Bruant *et al.*, 2010) was expressed in the *Acetobacterium* str. MES1 to convert acetyl-CoA to butyrate (aceto.peg.1850-4, 2312), which may explain the butyrate production by the electrosynthetic microbiome (Supplementary Figure 3)(Marshall *et al.*, 2013).

**Figure 3.**
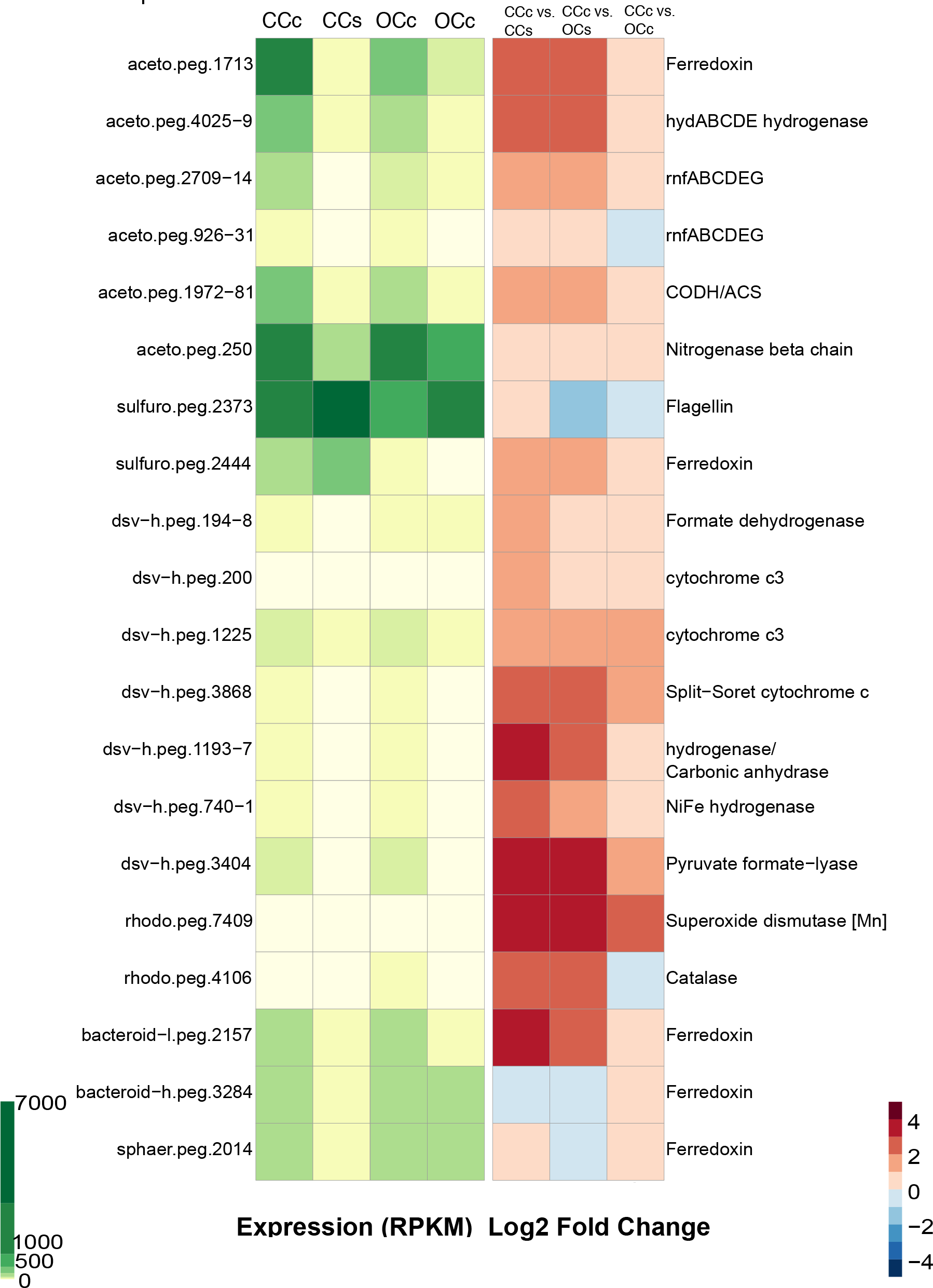
Expression profile and comparative expression of important genes. The green panel shows relative expression between microorganisms and the red panel compares differential expression between condition.

Reducing equivalents for the Wood-Ljungdahl pathway, reduced ferredoxin and NADH, were generated by a soluble uptake hydrogenase complex, *hydABCDE* (aceto.peg.4025-9), which were the most highly expressed *Acetobacterium* str. MES1-associated hydrogenase genes on the electrode. The high expression of *hydABCDE* and *rnfABCDEG* gene clusters, suggests *Acetobacterium* str. MES1 conserves energy similar to *A. woodii*.(Schuchmann and Müller, 2014) The exception is that MES1 has two Rnf complex copies, one with 15-fold greater gene expression on the electrode surface (high: aceto.peg.2709-14 and low: aceto.peg.926-931). This expression difference is similar to *Azotobacter vinelandii* whose copy expression disparity is driven by ammonium depletion.(Curatti *et al.*, 2005) Rnf expression positively correlates with nitrogen fixation (*nif*) gene expression, which is likely because Rnf is required for stable accumulation of the 4Fe4S clusters in *nifH*.(Jeong and Jouanneau, 2000; Curatti *et al.*, 2005) *Acetobacterium* str. MES1 maintains a nitrogenase gene cluster *(nifBEHN)* (aceto.peg.248-54) with an adjacent ferredoxin. In fact *nifH* was in the top 5 most expressed *Acetobacterium* str. MES1 genes in all conditions, suggesting N-limitation and/or an involvement in electron transfer (Figure 3). As ammonium is depleted, nitrogenase converts protons to hydrogen in the absence of N_2_ (Ryu *et al.*, 2014), which may contribute to the measured hydrogenesis. This supports previous observations of a lag time between fresh media exchange and the onset of hydrogenesis, and of a positive correlation between *Acetobacterium* and hydrogen production (LaBelle *et al.*, 2014). It is also possible that electron transfer from the Rnf complex to the nitrogenase for proton reduction is a source of hydrogen production by the electrosynthetic microbiome in this and related studies (LaBelle *et al.*, 2014), and is a potential target for biotechnological exploitation (Ryu *et al.*, 2014).

*Sulfurospirillum* str. MES7 remained consistently abundant (>60% relative abundance) in the supernatant despite frequent exchanges with fresh medium. However, the role of *Sulfurospirillum* in electrosynthesis has remained elusive (Marshall *et al.*, 2012, 2013). Given its prevalence it must fix CO_2_ and/or utilize acetate as a carbon source. In support of CO_2_ fixation, *Sulfurospirillum* spp. are hypothesized to fix CO_2_(Goris *et al.*, 2014) and other members of the epsilonproteobacteria can fix CO_2_ via the reductive tricarboxylic acid (rTCA) cycle (Hügler *et al.*, 2005). As shown here and a recent *Sulfurospirillum* comparative genomic study (Ross *et al.*, 2016), the *Sulfurospirillum* str. MES7 assembly contains several genes necessary to overcome the irreversible enzymes of the TCA cycle, including a 2-oxoglutarate oxidoreductase alpha, beta, delta, and gamma subunits (EC 1.2.7.3, sulfuro.peg.1237-40) and fumarate reductase (EC 1.3.5.1, sulfuro.peg.592-4). No ATP citrate lyase or citryl-CoA lyase were found, but it does have genes for the conversion of citrate to oxaloacetate + acetate (citrate lyase alpha and beta chains, and citrate (pro-3S)-lyase, EC 4.1.3.6, sulfuro.peg.2133-6), suggesting the possibility of CO_2_ fixation through rTCA. This is also supported by the transcriptional activity of key genes in the rTCA cycle (Supplementary Figure 4). In addition, acetate permease (sulfuro.peg.1580) and acetate kinase (sulfuro.peg.1274) are both highly expressed, indicating either that a supplementary organic carbon source might be required for growth, or that *Sulfurospirillum* str. MES7 flexibly switches between CO_2_ fixation and acetate oxidation in the MES reactor.

By far the most highly expressed genes relating to terminal electron accepting processes in *Sulfurospirillum* str. MES7 (for any condition) were cytochromes, particularly cytochrome c oxidases. C-type heme-copper oxidases are the final step in the catalytic reduction of oxygen in certain bacteria (Lee *et al.*, 2011) and *ccoNOPQ* is highly expressed by *Sulfurospirillum* str. MES7 (sulfuro.peg.1785-8) in all MES conditions. The high expression of *ccoNOPQ* suggests *Sulfurospirillum* str. MES7 is growing microaerobically in the MES, using oxygen as the terminal electron acceptor. This supports the hypothesis that microaerophilic bacteria provide a supportive role to the rest of the community by scrubbing low levels of oxygen that diffuse from the anode chamber.

*Desulfovibrio* str. MES5 was the 3^rd^ most abundant taxon, and likely converts CO_2_ to formate and while using cytochromes, formate dehydrogenase, and/or hydrogenases to accept electrons from the cathode. The high expression of formate dehydrogenase (3fold, dsv-h.peg.194) and cytochromes (6-fold, dsv-h.peg.1195; 6-fold, dsv-h.peg.3868) on the electrode compared to the supernatant supports this hypothesis. The lack of a suitable terminal electron acceptor indicates that *Desulfovibrio* str. MES5 reduces protons and generates hydrogen gas (Carepo *et al.*, 2002), which is supported by the comparatively high expression of hydrogenases (dsv-h.peg.740-1) on the electrode surface (>3-fold CCc vs. CCs). Furthermore, *Desulfovibrio* spp. have been shown to interact with a cathode to facilitate electrohydrogenesis.\ (Aulenta *et al.*, 2012; Croese *et al.*, 2011; Yu *et al.*, 2011). It is likely that small amounts of acetate from *Acetobacterium* str. MES1 are used as a supplementary carbon source for *Desulfovibrio* str. *MES5*, but based on the high hydrogenase and low acetate kinase expression pattern the majority of reducing potential comes from the electrode.

While *Acetobacterium* str. MES1, *Sulfurospirillum* str. MES7, and *Desulfovibrio* str. MES5 can explain most of the activity occurring in the microbial electrosynthesis systems, the remaining genomes have broad metabolic capabilities, but are typically in lower abundance (<7%). We hypothesize that they are consuming detritus, degradation products, and/or short chain fatty acids generated by the abundant organisms. A pangenome associated with the Rhodobacteraceae family had an expression profile consistent with acetate oxidation and cbb3-type cytochrome c oxidases that could be indicative of microaerobic growth. The latter have been expressed in *Rhodobacter* sp. growing in microaerobic and anaerobic conditions (Kaplan *et al.*, 2005) and were also found to be upregulated by *Shewanella* oneidensis MR-1 growing on electrodes (Rosenbaum *et al.*, 2012). While we are hypothesizing that these cytochrome c oxidases in Rhodobacteraceae str. MES12 and *Sulfurospirillum* str. MES7 are involved in microaerobic growth, given their prominence in this study and others involving bioelectrochemical systems (Rosenbaum *et al.*, 2012), their role in EET should be investigated further. One of two *Bacteroides* genomes (str. MES10) was closely related (93% sequence identity) to *Proteiniphilum acetatigenes*, which has been shown to generate acetate when growing on wastewater cell debris(Chen, 2005) and Croese et al. discovered a relatively high abundance of uncultured *Bacteroides* in a hydrogen producing biocathode community (Croese *et al.*, 2013), suggesting possible productive roles for the *Bacteroides* str. MES9 and MES10 in the electrosynthetic microbiome. Interestingly, *Bacteroides* str. MES9 expresses a butanol dehydrogenase (bacteroid-h.peg.1751) on the electrode surface, which suggests that changing the operating conditions of the MES could produce biofuels and other valuable products (see Bioprospecting section in supporting materials). Additionally, both *Bacteroides* genomes and the Sphaerochaeta str. MES8 genome exhibited high expression of ferredoxin genes (bacteroid-h.peg.3284, bacteroid-l.peg.2157, sphaer.peg.2014), which may be used to shuttle electrons for community metabolism.

### Metabolic model reconstructions and analysis

Genome-scale metabolic models were constructed for the three dominate species in the electrosynthetic community (see methods): *Acetobacterium* str. MES1, *Sulfurospirillum* str. MES7, and *Desulfovibrio* str. MES5. These models were then used, in combination with flux balance analysis (Orth *et al.*, 2010), to predict the metabolic activity of each of these species during microbial community growth on the electrode. The accuracy of these flux predictions was evaluated by calculating the fraction of active reactions predicted by the models that were associated with actively expressed genes, as determined from the metatranscriptomic data (Table 1 and methods). In this analysis, the *Acetobacterium* model displayed only one plausible flux profile, which involved carbon fixation and acetate production via the Wood-Ljungdahl pathway using hydrogen, raw electrons, or similar reducing equivalents as an electron source. With this flux profile, there were 338 active reactions with associated genes in *Acetobacterium*, and for 258 (76%) of these reactions, at least one of the associated genes was actively expressed. Most active reactions that lacked expression support were involved in either amino acid biosynthesis or nucleotide metabolisms. The *Sulfurospirillum* model predicted two alternative theories for the metabolic role of this species: (i) reduction of CO_2_ through the reductive TCA cycle with hydrogen used as the reducing agent (67% of active reactions associated with at least one expressed gene), and (ii) oxidation of acetate coupled to O_2_ reduction (66% of active reactions associated with at least one expressed gene). Given the nearly equal agreement of both of these operating with our transcriptome data, and given the high abundance of *Sulfurospirillum* in our community, it is possible *Sulfurospirillum* actually performs both roles depending on its context and environment. The *Desulfovibrio* model also predicted two alternative theories for the metabolic role of this species in our electrosynthetic microbiome: (i) conversion of CO_2_ and electrons/hydrogen to formate (75.7% of active reactions associated with at least one expressed gene), or (ii) consumption of acetate (75.9% of active reactions associated with at least one expressed gene). As with *Sulfurospirillum*, the expression data was equally supportive of both metabolic theories, indicating that *Desulfovibrio* also performs a combination of carbon-fixation and acetate utilization.

**Table 1.**
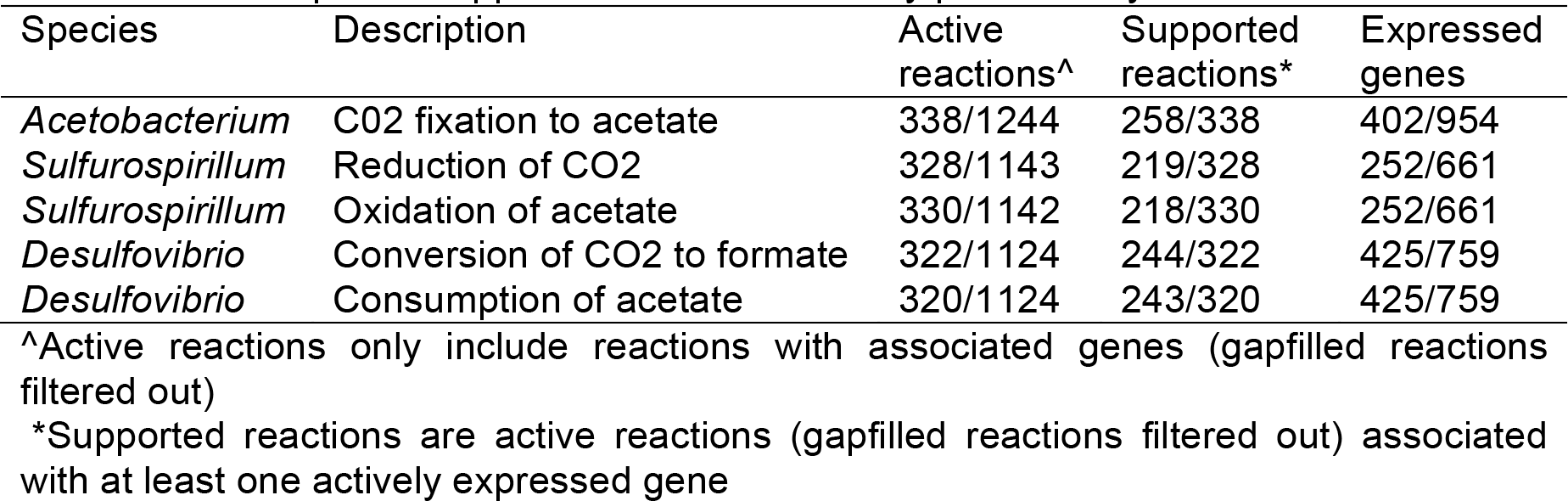
Transcriptome support for metabolic activity predicted by models

In addition to evaluating the overall evidence supporting each of our predicted flux profiles, the transcriptome data was also applied to evaluate the flux profiles on a pathway-by-pathway basis, enabling a better understanding of model accuracy, and revealing insights into potential interspecies interactions (see Methods and Figure 4). Generally, this analysis showed a large degree of agreement between predicted flux profiles and expression data, with some notable exceptions: (i) the terpenoid pathways in all our models were gap-filled because the models all include ubiquinone as a component of biomass, but there is little evidence for these pathways in our genomes and it is likely these genomes are functioning anaerobically and do not have to produce ubiquinone; (ii) the *Acetobacterium* and *Desulfovibrio* models both appear to overuse their pentose-phosphate-pathways compared to what would be expected based on expression data; (iii) the *Desulfovibrio* model overuses its thiamin pathway and should actually obtain thiamin from an external source according to expression data; (iv) the *Sulfurospirillum* model overuses its sulfur and nitrogen metabolism pathways; and (v) all three models appear to underutilize the vitamin B6 and fructose and mannose metabolism pathways.

**Figure 4.**
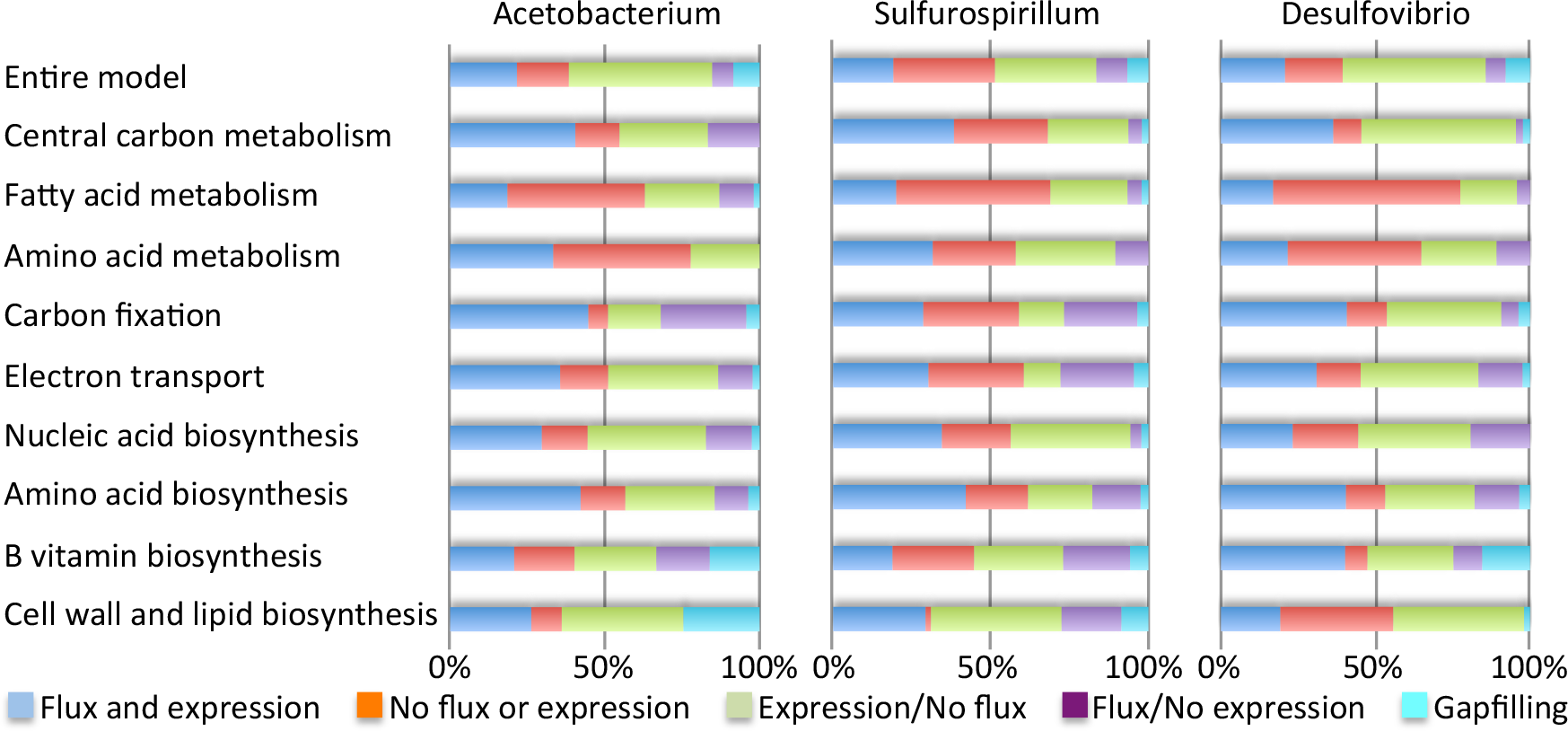
Pathway flux and model agreement with expression data. The degree of agreement between the model-based flux predictions and expression data for each of the three metabolic models is shown, both for the entire models and broken down by categories of metabolism. In the graph, reactions are divided into five categories based on their flux and the expression of their associated genes: (i) reactions that are active and associated with at least one expressed gene (dark blue); (ii) reactions that are inactive and associated only with unexpressed genes (dark red); (iii) reactions that are inactive and associated with one or more expressed genes (green); (iv) reactions that are active and associated only with unexpressed genes (purple); and (v) gapfilled reactions associated with no genes. The dark blue and dark red categories indicate agreement between the models and expression data; purple and green categories indicate disaareement.

The pathway-based model analysis (Figure 4) reveals interactions and dependencies among the top genomes. *Desulfovibrio* appears to require an external source of histidine, thiamin, riboflavin, and folate, while the other genomes could be sources of all four of these compounds due to depletion or absence in the supplied medium. *Acetobacterium* appears to be unique in making glutathione, and also has far more representation in carbon fixation than the other genomes. In contrast, *Sulfurospirillum* appears unique in possessing an active glyoxylate metabolism and the most active TCA cycle. It is interesting to note that all three genomes appear to produce most of their own amino acids, vitamins, and nucleotides, with *Desulfovibrio* being the least prototrophic of the three. Biochemical reaction equations, compounds, associated flux values, and comparative genomics tools including hypothetical gene knockout experiments for all the models can be found at the KBase Narrative Interface: https://narrative.kbase.us/narrative/ws.15248.obj.1.

## Discussion

Metabolic analysis and modeling of this unique microbial community capable of converting CO_2_ into volatile fatty acid provides a detailed look into how communities actively metabolize on an electrosynthetic biocathode. The genome-wide metabolic capabilities of microorganisms in the community were determined, and a putative metabolic model of the primary community members has been summarized (Figure 5). CO_2_ fixation and the carbon flux through the electrosynthetic microbiome center on the Wood-Ljungdahl pathway of the *Acetobacterium* genome, components of which were more active on the closed circuit electrode compared to any other condition tested, demonstrating the importance of electrode-associated growth by *Acetobacterium*. In addition, reducing equivalents in the form of hydrogen and reduced enzymes (eg. ferredoxin) also stem from the *Acetobacterium*, alongside major contributions from *Desulfovibrio* and *Sulfurospirillum*. Other organisms in the community are contributing to overall fitness by scrubbing toxic compounds (e.g. oxygen) and providing nutrient exchange. Given the proper conditions and applied drivers, this electrosynthetic microbial community has a wide range of biotechnological potential, including the production of alcohols, 2,3-butanediol, and polyhydroxyalkonoates (see supplementary note on bioprospecting and Supplementary Table 4 for more detailed information).

**Figure 5.**
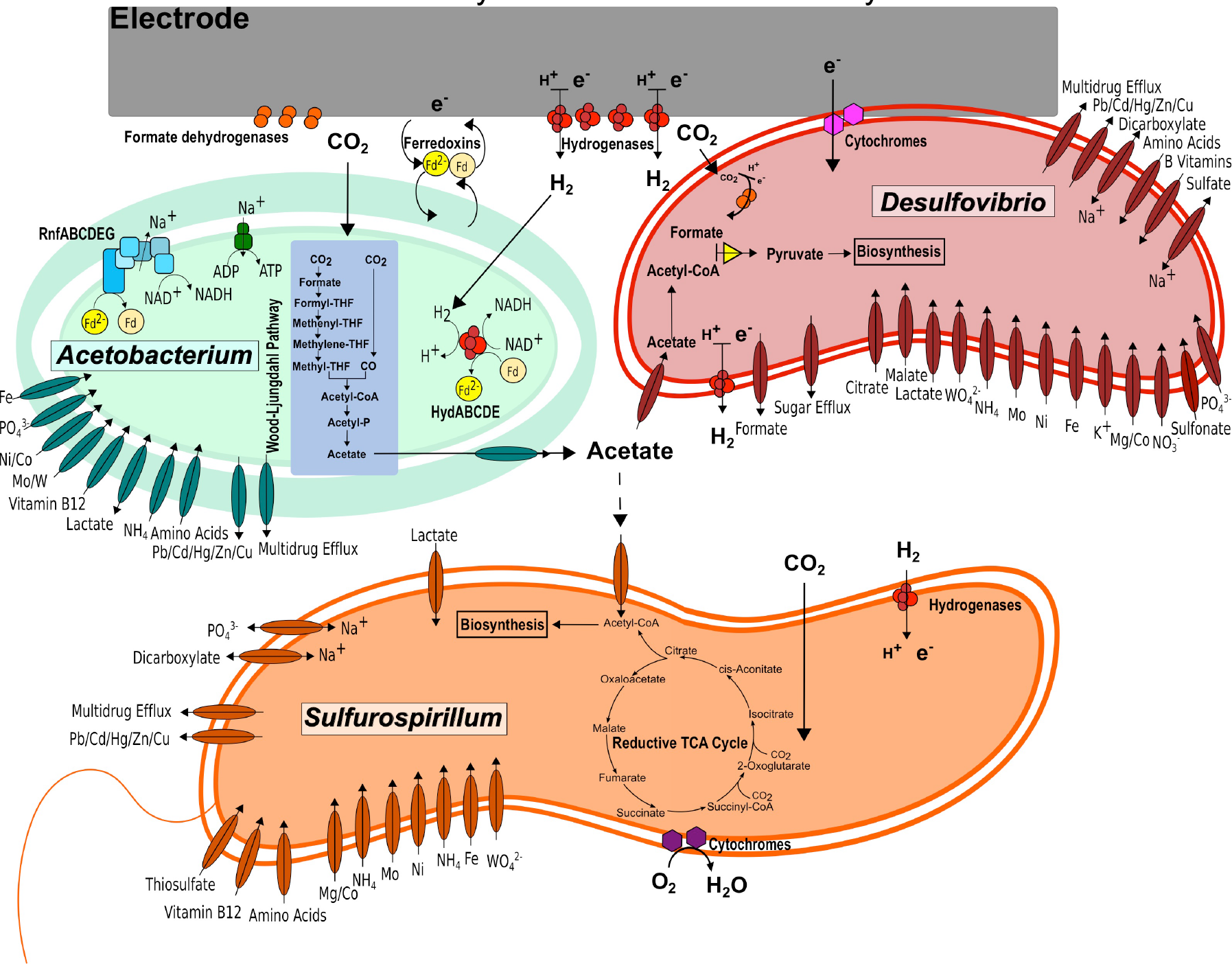
Hypothetical model of key metabolic activities and interactions among dominant members of the electrosynthetic microbial community.

Generally, the proposed mechanisms for cathodic electron transfer in the absence of added mediators are analogous to anodic electron transfer, namely that cytochromes are the primary conduit of electrons to and from the microbial cell (Holmes *et al.*, 2006; Leang *et al.*, 2010; Ross *et al.*, 2011; Inoue *et al.*, 2011; Carlson *et al.*, 2012). The role of cytochromes on anodes has been primarily elucidated in pure culture, but metatranscriptomic studies also point to cytochrome involvement in EET to anodes (Ishii *et al.*, 2013, 2015) and even less is known about EET in communities grown on cathodes. In this study, *Desulfovibrio* str. MES5 has three upregulated cytochromes on the electrode compared to the supernatant: two with a 6-fold increase (dsv-h.peg.1195 and dsv-h.peg.3868) and one with a 2.5-fold increase (dsv-h.peg.1225). The latter two were upregulated greater than 2-fold on the closed circuit cathode compared to the open circuit cathode. Additionally, a formate dehydrogenase from *Desulfovibrio* str. MES5 (dsv-h.peg.194) was upregulated nearly 3-fold on the electrode compared to supernatant, >2-fold CCc versus OCc, and could explain formate transiently observed in the supernatant (Supplementary Figure 3) (Marshall *et al.*, 2013)(da Silva *et al.*, 2013). In addition to their involvement in electron transfer at cathode surfaces, cytochromes can also act as natural mediators for hydrogenases (Pereira *et al.*, 1998; Yahata *et al.*, 2006) and could shuttle electrons from the electrode or from electrode-attached hydrogenases to the microbial community. Three NiFe hydrogenases were highly expressed by *Desulfovibrio* str. MES5 on the electrode compared to the supernatant (dsv-h.peg.740-741, >3-fold and 6-fold increase on electrode compared [CCc] to supernatant [CCs]; and dsv-h.peg.1193, 6-fold increase on electrode [CCc] compared to supernatant [CCs]). One of the NiFe hydrogenases (dsv-h.peg.1193) was adjacent to the upregulated high molecular weight cytochrome c (dsv-h.peg.1195). Given the high expression on the cathode compared to supernatant, it is hypothesized that cytochromes, hydrogenases, and possibly formate dehydrogenases associated with *Desulfovibrio* str. MES5 act as electron mediators between the electrode and the cell to deliver reducing equivalents. In addition to the well-established role of cell-bound cytochromes in direct electron transport with an electrode (Bond *et al.*, 2012; Pirbadian *et al.*, 2014), free cytochromes have been shown to be integral to anodic electron transfer while still imbedded in the biofilm matrix, and could be assisting electron transport to the community in this electrosynthetic biofilm (Inoue *et al.*, 2011).

The recent discovery that extracellular enzymes (likely hydrogenases and formate dehydrogenases) may be present in cathodic biofilms and may catalyze electron transfer into microbially-utilizable substrates (hydrogen and formate) adds a new and important piece to solve the EET mechanistic puzzle (Deutzmann *et al.*, 2015). Of the genes highly expressed on the electrode, several redox-active electron carriers (particularly hydrogenases and ferredoxins) had high differential expression on the poised electrode compared to the supernatant and open circuit electrode. A *Sulfurospirillum* str. MES7 ferredoxin (sulfuro.peg.2444) is >2-fold higher on the electrode compared to the supernatant (1.5x higher CCc vs. OCc) and the low abundance *Bacteroides str*. MES10 contains a ferredoxin (bacteroid-l.peg.2157) that is >9-fold upregulated on the electrode compared to supernatant (1.7x higher CCc vs. OCc). Importantly, a ferredoxin from *Acetobacterium* str. MES1 (aceto.peg.1713) was 7-fold higher on the electrode (CCc) compared to both supernatant conditions (CCs and OCs). Additionally, the highly expressed *Acetobacterium* str. MES1 hydrogenase gene cluster, *hydABCDE* (aceto.peg.4025-4029), had >5-fold higher expression per cell on the electrode compared to the supernatant (1.6x higher CCc vs. OCc), despite the presence of hydrogen in the supernatant. The combination of soluble hydrogenases and ferredoxins on the cathode could facilitate cathodic electron transfer into organisms like *Acetobacterium* that contain no cytochromes and no obvious means of electron transfer through the cell envelope.

Despite the lack of a redox-active signal in cell-free cyclic voltammograms and a catalytic wave characteristic of direct electron transfer in the electrosynthetic microbial community (Marshall *et al.*, 2013), SEM-EDX images show extracellular material with elemental signatures of common redox-active enzymes (Supplementary Figure 5). Additionally, no reduction in current or shift in the voltammetric peaks were observed when the supernatant was replaced with fresh medium, suggesting tightly bound electron transfer mechanisms. The discovery of metals such as iron and nickel on the electrode surface could be concentrated enzymes, but microbial or pH induced precipitation of the metals as (hydr)oxides, sulfides, carbonates, phosphates, perhaps even as nanoparticles cannot be ruled out. The latter has been hypothesized by Jourdin et al. who observed copper concentrated on the surface of a microbial electroacetogenic cathode (Jourdin *et al.*, 2016).

Based on the highly expressed redox active proteins on the electrode (all of the proteins mentioned above are in the top 5% of total genes expressed on the electrode), and the consistent hydrogen production in the reactors (Marshall *et al.*, 2012, 2013; LaBelle *et al.*, 2014), extracellular enzymes and/or concentrated metals are likely functionalizing the electrode while hydrogen, ferredoxins, and/or cytochromes act as shuttles for different organisms in the biofilm. This would explain why microbes lacking an outer membrane can thrive on an electrode, and why previous studies ran electrohydrogenic biocathodes that could sustain hydrogen generation in the absence of a carbon source for over 1000 hours (Rozendal *et al.*, 2008). Further studies are underway to fully characterize functionalization of electrodes by microbes so as to optimize this strategy.

The modeling results demonstrate how metabolic models are useful for interpreting expression data in order to predict the role of individual species participating within a mixed microbial community. The model predictions: (i) confirmed the role of *Acetobacterium* as the primary species utilizing electrons to reduce CO2; (ii) identified the more complex behavior of *Sulfurospirillum* as both reducing CO2 and utilizing acetate; and (iii) provided more depth and detail into our understanding of the heterotrophic growth of *Desulfovibrio*. These models refined models should prove to be a resource for ongoing efforts to understand, and ultimately design and refine electrosynthetic microbiomes.

We have provided insights into the genetic basis of electrosynthetic microbial communities, which can be used for the optimization of microbial electrosynthesis through operational changes and eventual pathway engineering. Furthermore, it was demonstrated that a diverse set of microorganisms could be active in limited niche space with carbon dioxide as the only carbon source and the electrode as the only electron donor. The discovery that predominant members of the community provide an ecosystem service by scrubbing oxygen makes this, and similar, electrosynthetic microbial communities valuable for practical application. If deployed in true carbon-capture situations such as power plants or industrial exhaust lines, oxygen scrubbing to protect the electrosynthetic anaerobes will be important. Finally, the comprehensive analysis of the genomes and development of metabolic models provide the framework to boost production rates and elevate this system to a platform chemical synthesis technology.

## Acknowledgements

We thank Drew Latta for SEM-EDX work and Fangfang Xia and Sebastien Boisvert for initial metagenome assembly advice. C.W.M. was supported in part by an Argonne National Laboratory Director's Fellowship. The project was funded by the Department of Energy, Advanced Projects Research Agency-Energy (DE-AR0000089) and Office of Naval Research: Grant# N00014-15-1-2219. We also acknowledge the University of Chicago Research Computing Center for their support.

